# Scalable metabolic pathway analysis

**DOI:** 10.1101/2020.07.31.230177

**Authors:** Ove Øyås, Jörg Stelling

## Abstract

The scope of application of genome-scale constraint-based models (CBMs) of metabolic networks rapidly expands toward multicellular systems. However, comprehensive analysis of CBMs through metabolic pathway analysis remains a major computational challenge because pathway numbers grow combinatorially with model sizes. Here, we define the minimal pathways (MPs) of a metabolic (sub)network as a subset of its elementary flux vectors. We enumerate or sample them efficiently using iterative minimization and a simple graph representation of MPs. These methods outperform the state of the art and they allow scalable pathway analysis for microbial and mammalian CBMs. Sampling random MPs from *Escherichia coli*’s central carbon metabolism in the context of a genome-scale CBM improves predictions of gene importance, and enumerating all minimal exchanges in a host-microbe model of the human gut predicts exchanges of metabolites associated with host-microbiota homeostasis and human health. MPs thereby open up new possibilities for the detailed analysis of large-scale metabolic networks.

## Introduction

Constraint-based models (CBMs) can represent the complete genome-scale metabolic network of an organism and allow for the analysis of steady-state fluxes (reaction rates) based only on stoichiometry and other physico-chemical constraints. Large collections of curated models^1,2^ and tools for automated model generation^3^ make CBMs available for a wide range of organisms. Their scope of application is also expanding from unicellular to multicellular systems such as microbial communities or human tissues, increasing the need for analysis methods that scale to large metabolic networks^4^.

The common analysis of CBMs assumes that metabolism is essentially at steady state on the time scale of other cellular processes. This produces a linear system of *m* metabolite mass balances in which the variables are the *n* fluxes **r** ∈ ℝ^*n*^ constrained by the stoichiometric matrix **N** ∈ ℝ^*m×n*^,

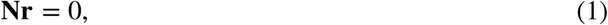

and any other linear constraints, e.g., upper and lower flux bounds:

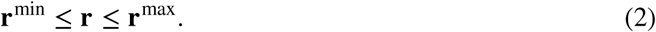

Solutions that satisfy these constraints are feasible combinations of steady-state fluxes known as flux distributions^5^.

The most elementary method for analyzing flux distributions is flux balance analysis (FBA), in which an assumed objective is maximized or minimized subject to constraints (1) and (2) in a linear program (LP):

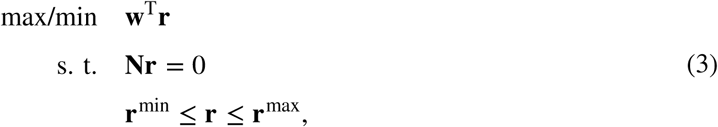

where **w** is the vector of weights for the fluxes. Maximal growth rate, as represented by the flux of a biomass reaction, is usually assumed as the objective for microbes^6^. FBA scales to very large networks thanks to efficient LP solvers, but it only returns a single optimal solution that is not necessarily unique^7^.

In contrast to FBA and other biased methods, unbiased approaches can characterize the whole solution space. Random flux sampling can be used to obtain probability distributions over fluxes, but uniform sampling in high-dimensional spaces is challenging^8^. Pathway analysis is more comprehensive, defining all possible solutions in terms of sets of pathways such as elementary flux modes (EFMs)^9^ or elementary flux vectors (EFVs)^10,11^, as illustrated in Fig. 1. However, the number of pathways grows combinatorially with network size^12,13^. Hence, both approaches pose formidable challenges for large-scale analysis of metabolic networks.

**Figure 1.**
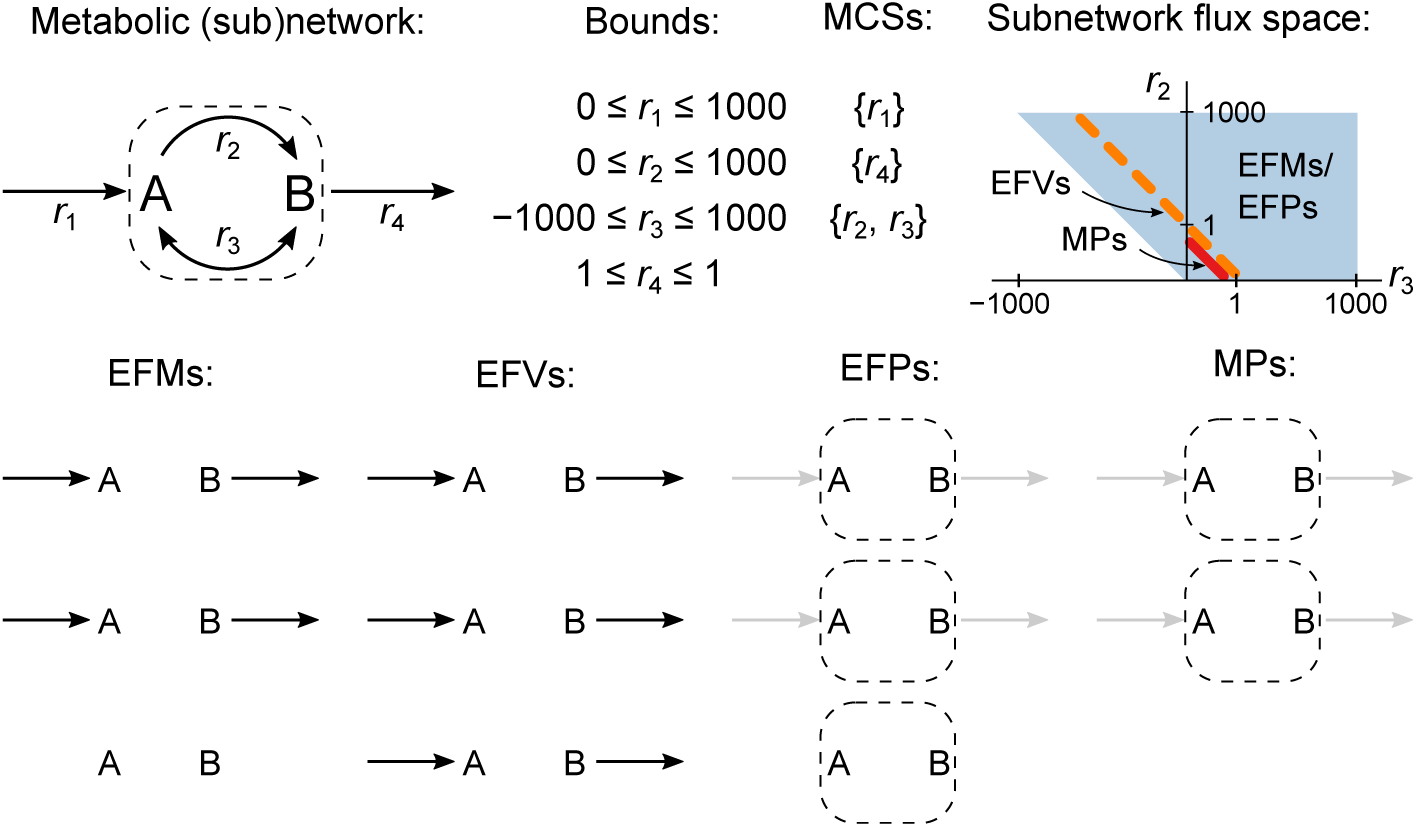
Example network with comparison of EFMs, EFVs, EFPs, and MPs. Two metabolites, A and B, participate in four reactions, *r*_1_–*r*_4_. The boundary reactions *r*_1_ and *r*_4_ exchange A and B with the environment and the internal reactions *r*_2_ and *r*_3_ define a subnetwork. All reactions except *r*_3_ are irreversible, and we apply the maximum bound ± 1,000 to all fluxes except *r*_4_, which is fixed to 1. The pathways defined by the EFMs, EFVs, EFPs, and MPs of the (sub)network are shown below. Reactions that are not active in a pathway are not shown and active reactions that are not in the subnetwork are shown in gray for EFPs and MPs. EFMs and EFVs can be used to find all the MCSs of the network, and EFPs and MPs can be used to find all MCSs of the subnetwork, in this case {*r*_2_; *r*_3_}. The pathways span different subnetwork flux spaces: EFMs and EFPs (blue) do not take bounds into account and their flux space is unbounded in the direction of positive *r*_2_ and *r*_3_, while EFVs (dashed orange line) and MPs (red line) account for bounds and span bounded subspaces of the EFM/EFP space. There are fewer MPs than other pathways and the MPs are the only pathways that do not include loops (*r*_3_ < 0).

Many conceptual and computational advances have been made to address these challenges. State-of-the-art algorithms can enumerate EFMs in networks with hundreds of reactions^14^, and modularization enables enumeration of billions of EFMs^15^. To circumvent combinatorial explosion and to obtain sets of pathways amenable to analysis, one can limit enumeration to a subset of pathways^16^. Examples of such subsets are the shortest EFMs^17^, random EFMs^18^, and EFMs that are consistent with thermodynamics^19^, regulation^20^, or metabolomics data^21^. Elementary flux patterns (EFPs), defined as the parts of EFMs in a network that pass through a subnetwork (Fig. 1), have also been introduced^22^. However, while these approaches reduce the number of pathways, they do not scale to large networks.

EFMs are also limited because they are only defined for homogeneous constraints such as (1) or reaction reversibilities. Elementary flux vectors (EFVs) generalize EFMs by allowing inhomogeneous constraints such as (2) or any other linear constraint, but the challenges of enumeration remain^10,11^. Alternative approaches aim to find minimal sets of reactions^23–25^ or precursor metabolites^26,27^ under inhomogeneous constraints. However, they all rely on direct minimization of the same mixed-integer linear program (MILP), which prevents scaling to large networks. The same MILP is also used to compute minimal cut sets (MCSs), minimal sets of reactions that, when inactivated, block specific network functions^28^.

Here, we introduce minimal pathways (MPs) as a concept well-suited for scalable pathway analysis (Fig. 1), and we present new methods for finding, sampling, and enumerating them in large metabolic (sub)networks. Our enumeration algorithm builds on recent work showing that the MILP can be split into an LP and a binary integer program (BIP), which can be alternated to compute EFMs and MCSs sequentially^29^. Our methods outperform the MILP approach when applied to a range of microbial and mammalian genome-scale CBMs, and we demonstrate applications to the analysis of central carbon metabolism in a genome-scale context and of host-microbe interactions in the human gut.

## Results

### Minimal pathways and detection methods

We define an MP as a support-minimal subset of reactions from a metabolic subnetwork, which in turn is a subset of reactions from a full metabolic network. Hence, all reactions in an MP need to be active (have non-zero flux) when no other reactions in the subnetwork are active in order to satisfy all constraints on the network as a whole. Combinations of fluxes in the subnetwork that satisfy all constraints on the full network lie in the projection of the solution space of the full network onto the subspace defined by the subnetwork (Fig. 1). The minimal set of vectors that generate this space without cancellations are its EFVs^10,11^, and the subset of EFVs that are support-minimal is equivalent to the set of all MPs for a given subnetwork and network.

To find MPs efficiently, we developed a novel iterative minimization approach. It leverages support-minimality: to find an MP, we iteratively remove reactions from the subnetwork and minimize subnetwork flux with an LP until no further reactions can be removed without disrupting the functional requirements of the network. A randomized version allows random sampling of MPs. To enumerate MPs and MCSs, we alternate iterative minimization with computation of cut sets in a separate BIP, as recently suggested for EFMs^29^. We also realized that maximal cliques in a simple graph representation of known MPs can predict subnetworks likely to contain unknown MPs, thereby reducing the sizes of optimization problems during enumeration.

To illustrate the different approaches, we consider MP enumeration and sampling in a random subnetwork of 562 reactions in the genome-scale *E. coli* model iML1515^30^ with 2,868 irreversible reactions. Here, graph-based ennumeration found all 27,648 MPs within 45 minutes (Fig. 2A), but after one hour runtime, the other methods had found an order of magnitude fewer MPs (Fig. 2B). The graph-based approach also found all MCSs, primarily at iterations where the full subnetwork was used rather than a subset defined by a clique from the graph (Fig. 2A,C). Notably, iterative minimization without graph found all MCSs very quickly, while the graph-based approach found them more gradually (Fig. 2C). In more detail, direct minimization only identified MPs of the minimal size 14, iterative minimization without graph and randomization found MPs in arbitrary but clearly not random order, and randomized iterative minimization found MPs in apparently random order (Fig. 2D). The order of MP sizes for graph-based enumeration was more structured than for the other iterative approaches; there was a tendency to first find small MPs and to find many MPs of similar size after one another (Fig. 2D). The effective subnetwork size in graph-based enumeration varied between the full subnetwork when no new cliques were available from the graph and much smaller sizes approximately equal to the sizes of MPs when cliques were available (Fig. 2E). Specifically, the cliques from the graph corresponded exactly to MPs in 70% of the cases, and deviated only by one reaction from the MP otherwise. For the other methods, subnetwork size was constant and equal to the size of the full subnetwork.

**Figure 2.**
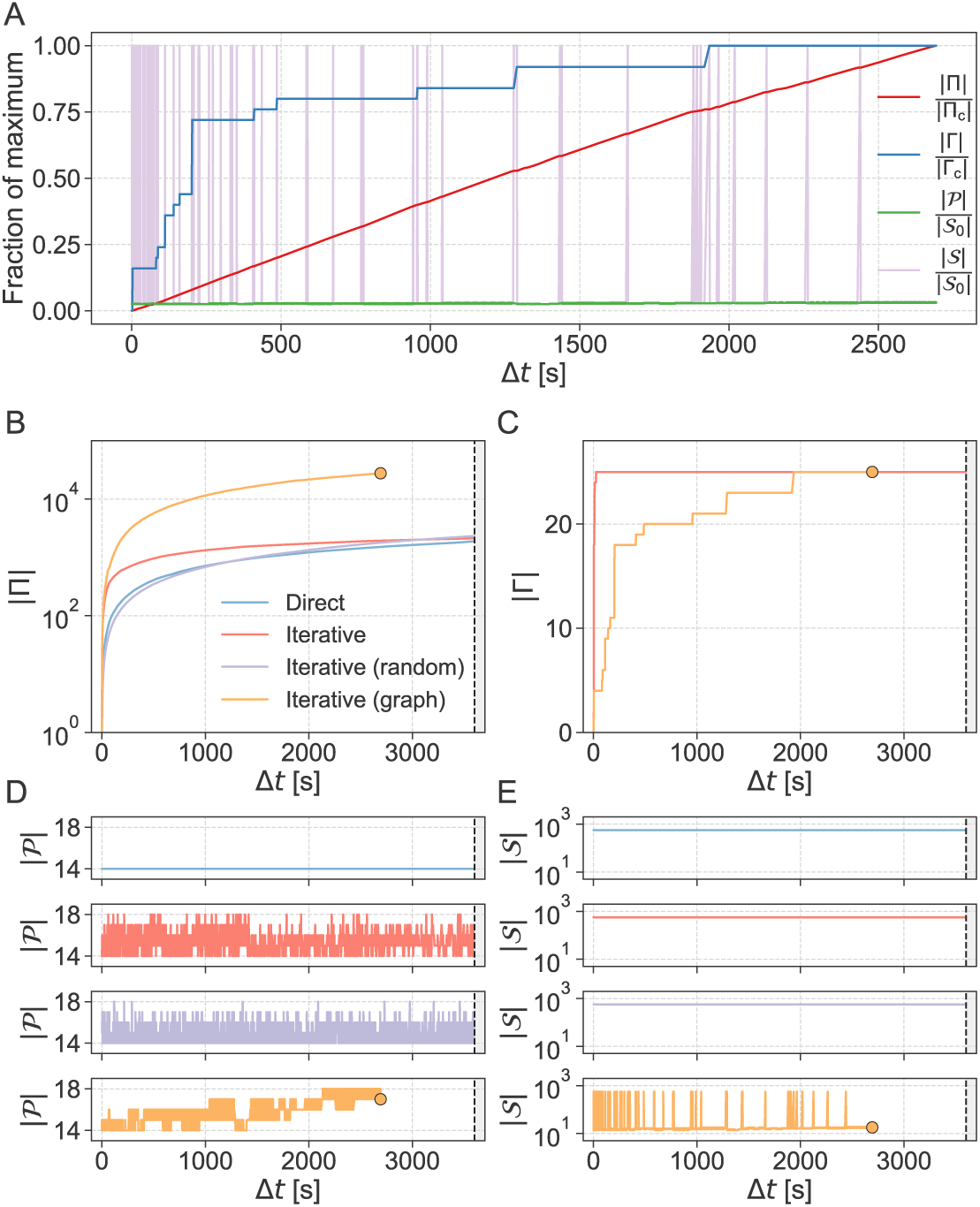
Example enumeration with comparison of methods. (**A**) Complete enumeration of MPs and MCSs using iterative minimization with graph in a random subnetwork of size |𝒮_0_| = 562 within the genome-scale *E. coli* model iML1515 with 2,868 irreversible reactions. The number of MPs |Π| and MCSs |Γ| found is shown along with the MP size |𝒫| and effective subnetwork size |𝒮| as a function of running time Δ*t*. Values are normalized by the size of the complete set of MPs |Π_c_|, the size of the complete set of MCSs |Γ_c_|, or the initial subnetwork size |𝒮_0_|. (**B-D**) Comparison to incomplete enumerations in the same subnetwork performed with direct minimization and iterative minimization with and without randomization, shown separately for |Π| (**B**), |Γ| (**C**), |𝒫| (**D**), and |𝒮| (**E**). End points of the completed enumeration are indicated by dots.

### Benchmarking of methods

We benchmarked our algorithms using six models of microbial and mammalian cells, ranging in size from about 100 to more than 14,000 irreversible reactions (Table S1). Specifically, we sampled 100 subnetworks each of sizes ranging from 10 reactions to the full network (**Supplementary Data 1** and **Methods**). The running time needed to find the first MP was always greater and grew much faster with subnetwork size for direct than for iterative minimization, demonstrating that our method is faster and scales to larger networks than the state of the art (Fig. 3A,B and Fig. S1). Direct minimization failed to find a single MP in many of the largest subnetworks across all models except the smallest model of *E. coli* core metabolism (Fig. S2A). Randomized iterative minimization outperformed direct minimization across all subnetwork sizes in smaller models and the largest subnetworks in larger models, but it was slower for intermediate subnetwork sizes in larger models (Fig. 3A,C). In addition to subnetwork size, running time should depend on the sizes of the full networks in which the subnetworks were embedded.

**Figure 3.**
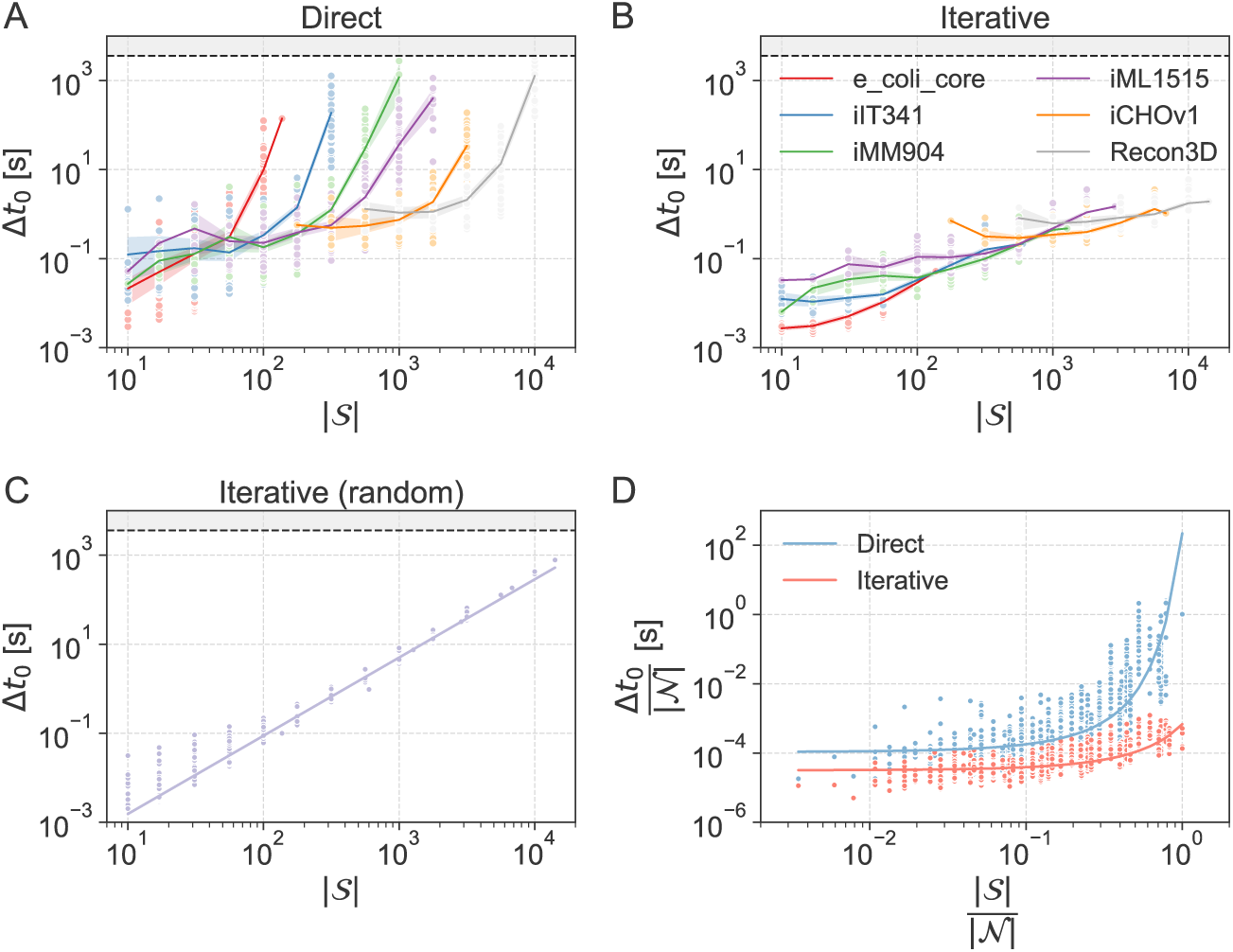
Running times for finding the first MP. (**A**)-(**C**) Running time *t*_0_ as a function of subnetwork size |𝒮| for direct minimization (**A**), iterative minimization (**B**), and randomized iterative minimization (**C**). Lines indicate means with 95% confidence intervals from bootstrapping and colors indicate models. Enumeration was stopped after one hour (indicated by dashed line). (**D**) Running time *t*_0_ as a function of subnetwork size |𝒮|, both normalized by network size |𝒩|, for direct (blue) and iterative (red) minimization. The double exponential model log *y* = *a* + *b*^*x*^ was fitted, yielding parameter estimates of *a* = −4.97 ± 0.02 and *b* = 7.32 ± 0.13 for direct minimization and *a* = −5.50 ± 0.01 and *b* = 2.33 ± 0.02 for iterative minimization without randomization.

Indeed, normalizing running time and subnetwork size by the size of the full network led to similar scaling across all models for direct minimization and iterative minimization without randomization according to a double exponential model (Fig. 3D). However, the running time of randomized iterative minimization scaled with absolute subnetwork size, following a monomial model (Fig. 3C).

Our graph-based approach completed more enumerations than any other method across all models, including all enumerations that were completed by any of the other methods (Fig. S2B). Reflecting the double exponential scaling of running time, the number of MPs found in enumerations that were completed within one hour grew superexponentially with subnetwork size (Fig. S3A). This was also the case for the number of MCSs obtained from enumerations using iterative minimization, but the set of all MCSs was generally smaller than the set of all MPs for subnetworks from all models except the (small) *E. coli* core model (Fig. S3B).

For enumerations that were not completed within one hour, the methods clearly differed in the number of MPs found. For direct minimization, the number of MPs decreased rapidly with subnetwork size for all models (Fig. 4A and Fig. S3). In contrast, iterative minimization without graph or randomization found approximately the same number of MPs (on the order of 10^3^) in all subnetworks across all models (Fig. 4B). For randomized iterative minimization, MP numbers in incomplete enumerations increased with subnetwork size until a model-dependent critical size was reached, after which they decreased linearly with little variation between subnetworks and models (Fig. 4C)). In most subnetworks, an order of magnitude more MPs were found by iterative minimization with graph than by any other method, but the number of MPs decreased more with subnetwork size when a graph was used (Fig. 4B,D). More MPs were found with randomized than with graph-based iterative minimization in some smaller subnetworks, but scaling was much worse with randomization (Fig. 4C,D).

**Figure 4.**
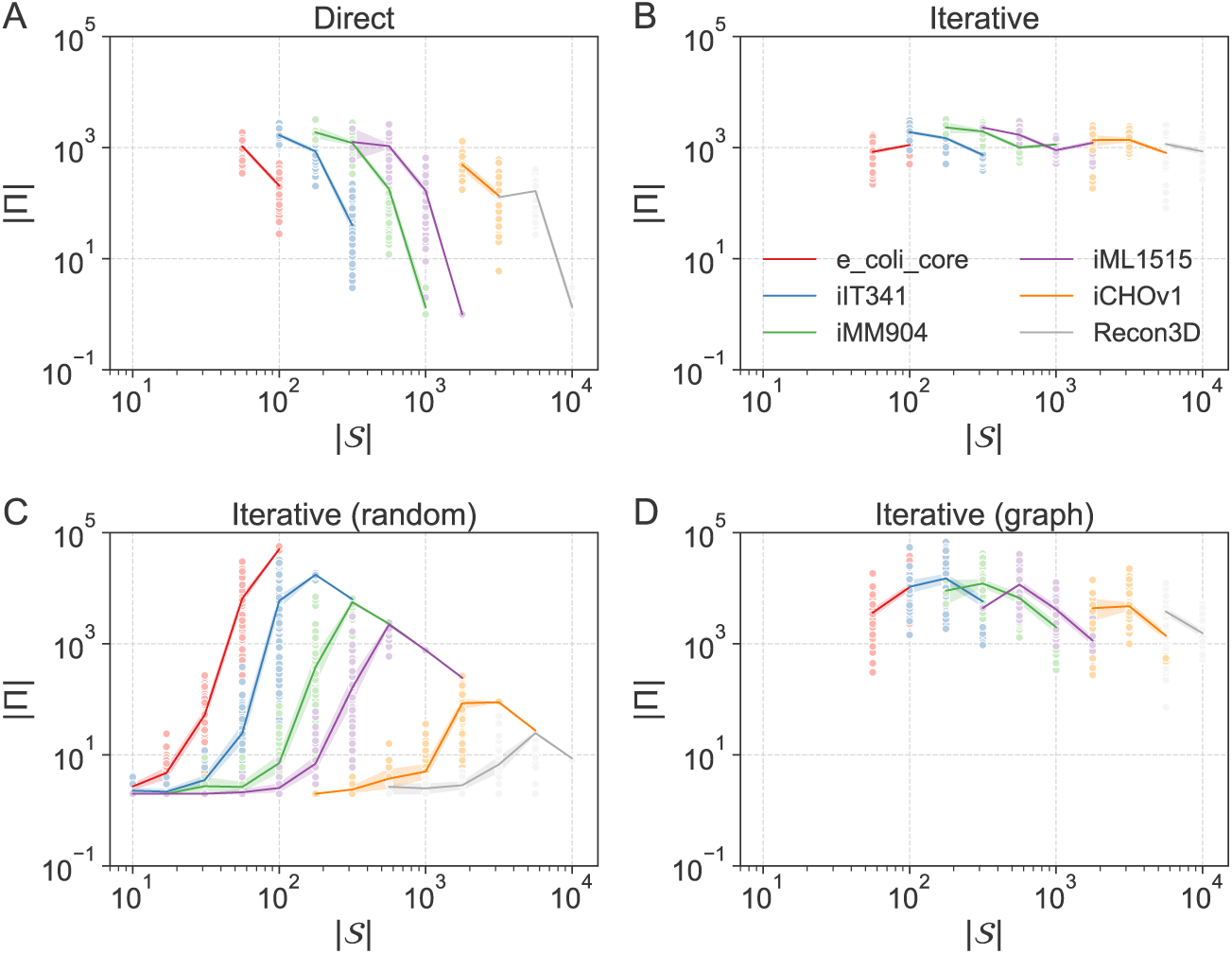
MPs found in incomplete enumerations. Number of MPs |Π| found in one hour as a function of subnetwork size |𝒮| for (**A**) direct minimization, (**B**) iterative minimization, (**C**) randomized iterative minimization, (**D**) and iterative minimization with graph. We excluded enumerations that were completed within one hour, except for randomized iterative minimization, where completion cannot be detected.

Thus, iterative minimization was systematically faster than direct minimization and allowed scaling all the way to the full networks. Randomization allowed scalable random sampling and our graph-based approach accelerated enumeration.

### Gene essentiality in central metabolism

To demonstrate the usefulness of MP sampling, we first analyzed gene essentiality, that is, whether a gene is required for the organism’s survival and reproduction. This is not only a standard test scenario for CBMs, but gene essentiality is also increasingly recognized as quantitative and context-dependent^31^. Even for model organisms such as *E. coli* it still unclear what the essential genome is, with targeted knock-out libraries^32^, randomized transposon mutagenesis^33^, and most recently CRISPRi screens^34,35^ yielding consensus as well as divergent gene sets.

We applied our randomized iterative minimization procedure to the central metabolism of *E. coli* as a subnetwork in the genome-scale model iJO1366^36^. This subnetwork of 153 irreversible reactions, mostly from glycolysis, the tricarboxylic acid (TCA) cycle, and the pentose phosphate pathway, is embedded in the full model of 3,219 irreversible reactions. Specifically, we sampled 100,000 MPs (in 10.3 h with 0.37 ± 0.18 s per iteration) for aerobic growth on a minimal glucose medium.

To reveal the relative importance of reactions and enzymes in central carbon metabolism, we calculated the fraction of MPs in which each reaction or enzyme participates (Fig. 5A, **Supplementary Data 2 and 3**, and **Methods**). No enzymes in the pentose phosphate pathway were essential, but five enzymes participated in more than 80% of all MPs (transaldolase, transketolases, ribulose 5-phosphate 3-epimerase, and ribose-5-phosphate isomerase). Also, four enzymes stood out in the TCA cycle (citrate synthase, aconitase, isocitrate dehydrogenase, and fumarase), appearing to be much more important for aerobic growth on glucose than, for example, 2-oxogluterate dehydrogenase and succinyl-CoA synthetase. As expected, isozymes such as the malate dehydrogenases in the TCA cycle were of very similar and intermediate importance. These patterns are consistent with the alternative operation of the full TCA cycle and the PEP-glyoxylate cycle for complete glucose oxidization^37^.

**Figure 5.**
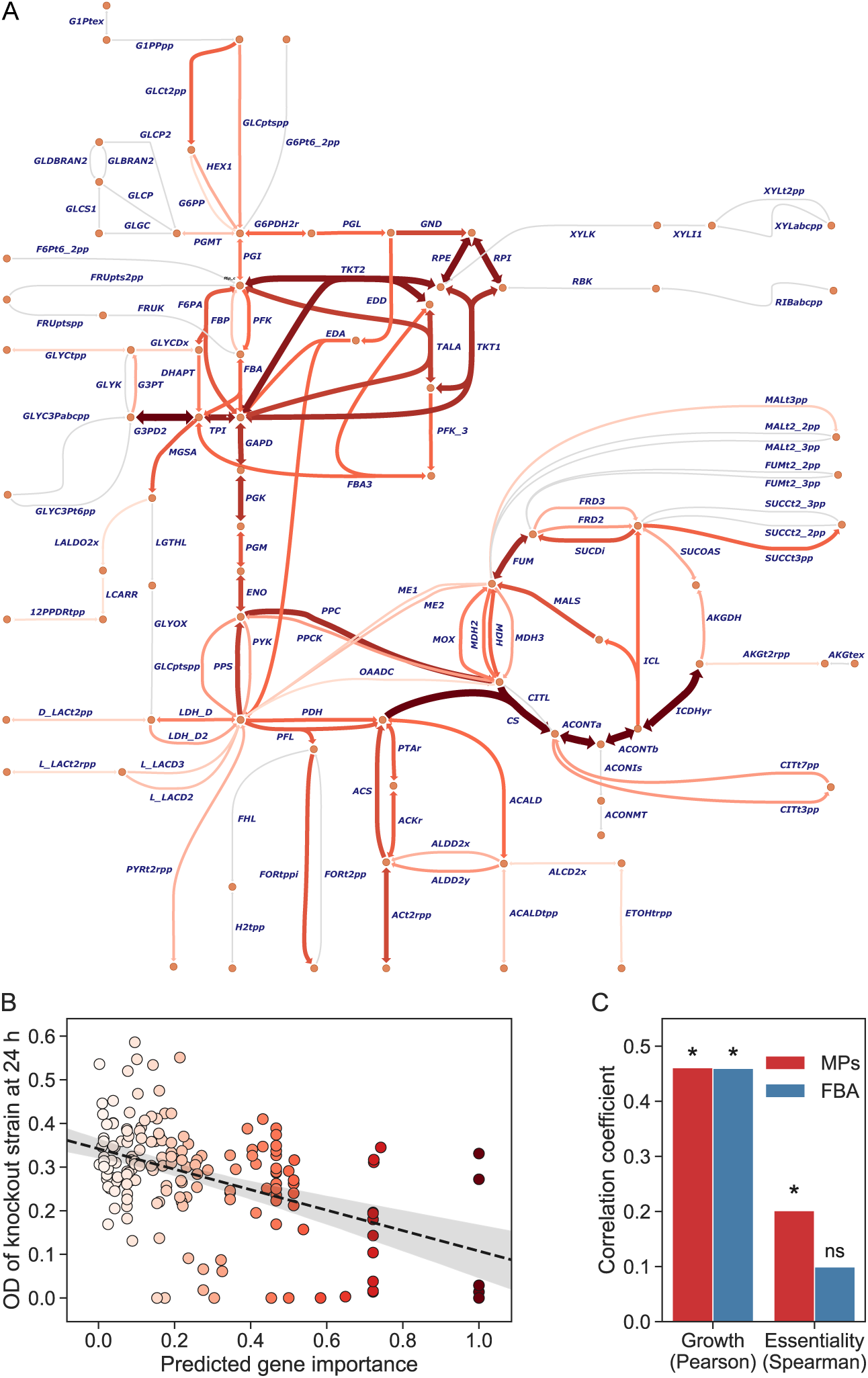
Example application of MP sampling. (**A**) Pathway map of *E. coli* central carbon metabolism. Circles are metabolites and arrows are reactions. The color and width of arrows indicate the fraction of 100,000 randomly sampled MPs in which reactions participated. The network was used as a subnetwork within the genome-scale model iJO1366 with a growth requirement of 0.1 h^−1^, a minimal glucose medium, and aerobic conditions. For reversible reactions, the fraction includes all MPs with the reaction regardless of flux direction. (**B**) Optical density (OD) of gene knockout strain at 24 h as a function of gene importance predicted by MP sampling (also indicated by color). A fitted line is shown with 95% confidence interval from bootstrapping. (**C**) Absolute correlation of gene importance predicted by MP sampling (red) and FBA (blue) with growth (OD of knockout strain at 24 h) and experimental essentiality score. Stars indicate significance (*p* < 0.05).

To evaluate our predictions quantitatively, we next mapped samples to enzyme-encoding genes and compared the relative importance of the genes to available growth and gene essentiality data from the Keio collection^32^. Gene importance in terms of growth reduction upon gene deletions predicted by MP sampling of central carbon metabolism correlated as well with growth of knock-out strains as FBA predictions for the full model (Fig. 5B and Fig. S5; Pearson’s *r* = −0.46, *p* = 2 ⋅ 10^−10^ for MPs). Also, both approaches yielded identical results for the binary classification of essential and non-essential reactions based on knock-out strains. In a previous elementary mode analysis with reduced models, however, small deviations in predictions were observed^38^. Finally, we analyzed the methods’ power of predicting Keio essentiality scores; they result from aggregating targeted knock-out and transposon mutagenesis data sets and range from −4 (indicating non-essentiality) to +3 (indicating essentiality). MP sampling clearly outperformed growth predictions from FBA in recapitulating these measures of experimental gene essentiality (Fig. 5C and Fig. S5), indicating that MP analysis may better reflect context-dependency in the data than point estimates obtained by FBA.

### Metabolite exchanges in the human gut

To show that our graph-based approach enables complete enumeration of MPs in large models of multicellular systems, we applied it to a host-microbe model of the human gut. We know little about how different members of the gut microbiota interact with each other and with the host, in particular with respect to the metabolite exchanges involved^39^. However, experimental studies have shown that metabolic pathway-rather than species-associations are predictive of host-microbe metabolic interactions^40,41^, making pathway analysis a suitable approach. Yet, whereas computational tools for the analysis of larger microbial communities have been developed (mostly as variants of FBA), applications are missing so far in general^39^, and no comprehensive pathway analysis exists specifically.

In our host-microbe model (see **Methods** for details), a human cell interacts with six commensal microbial species. We required ATP production for the human cell and growth for the microbial cells in anaerobic conditions and a complex metabolic environment. We used our recent approach to constrain the model to explain as much of the functional requirements as possible in terms of metabolite exchanges^42^, and enumerated all minimal sets of metabolite exchanges between microbes and from microbes to the human cell using iterative minimization with graph.

In this subnetwork of 3,209 metabolite exchange reactions within the full network of 33,291 irreversible reactions, we found all 240 MPs and 63 MCSs in 0.52 h with 7.6 ± 3.3 s per iteration (Fig. 6A). Because we constrained the problem to functionally required exchanges for given growth conditions, the MP set was small, indicating a focus on important exchanges (relaxations yield larger sets of MPs, but this was not the focus of our study). More specifically, the MPs were different combinations of 17 uptakes and 18 secretions, corresponding to 28 unique pairwise exchanges of ten different metabolites: amino acids, fermentation products, TCA intermediates, and the B vitamin niacin (Fig. 6B). The human cell was predicted to consume microbial formate, glycine, glycerol, malate, and ornithine, and the microbes exchanged all metabolites except formate and glycerol.

**Figure 6.**
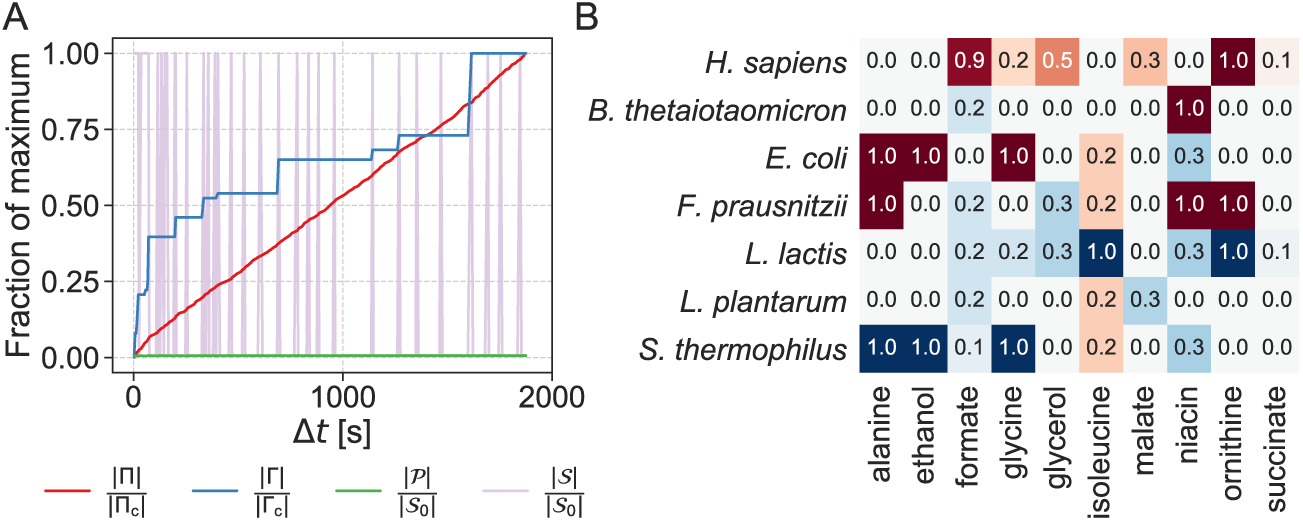
Example application of MP enumeration. (**A**) Complete enumeration of MPs and MCSs using iterative minimization with graph in a subnetwork of 3,209 metabolite exchanges within a host-microbe model with 33,291 irreversible reactions (symbols used as in **Fig. 2A**). (**B**) Fraction of MPs with uptake (red) and secretion (blue) for each metabolite (columns) and organism (rows).

Importantly, the metabolites predicted to be exchanged have been observed in the gut and implicated in the maintenance of host-microbiota homeostasis as well as human health and disease. For example, a possible alanine exchange predicted by the same microbial model collection used here was confirmed *in vitro*^43^, increased isoleucine levels have been associated with improved growth in humans^44^, and consumption of glycine by the microbiota has been shown to reduce glycine levels in the host, which has been associated with non-alcoholic fatty liver disease (NAFLD), obesity, and type 2 diabetes^45^. The gut microbiota also contributes more directly to NAFLD by producing ethanol, which suggests that consumption of ethanol through cross-feeding could be important in healthy gut microbiotas^46^. Glycerol degradation pathways are enriched in gut microbiotas of people with functional constipation^47^, elevated levels and cross-feeding of microbially derived formate have been associated with inflammation^48^, and levels of succinate as well as microbes producing or consuming it have been associated with obesity and related diseases^49^. Exchange of niacin (B3) and its importance for microbiota stability are supported by *in silico*, *in vitro*, and *in vivo* evidence^40,50,51^, and microbial ornithine production promotes healthy gut mucosa through crosstalk with the host^52^. Thus, the predicted exchanges appear plausible, and they amount to hypotheses on specific producers and consumers, which are hard to identify experimentally.

## Discussion

Our definition of MPs in terms of EFVs makes it straight-forward to reconcile MPs with available theory and tools, while providing several advantages over existing pathway definitions. First, as EFVs, MPs account for all linear constraints on the network as a whole, including inhomogeneous constraints neglected by EFMs. However, because MPs are a subset of the EFVs, there are fewer MPs than EFVs to enumerate. Second, all reactions in an MP contribute to fulfilling functional requirements as defined by constraints on the full network, which is not the case for EFMs, EFPs, or all EFVs. While this implies that MPs do not account for all possible flux distributions of the subnetwork, they cover the minimal flux distributions that support function, which have been shown to be consistent with omics data^53,54^. Also, MPs avoid the pervasive problem of thermodynamically infeasible loops^55^, which cannot contribute to well-defined functional requirements. Third, like other pathway definitions, MPs can be used to find all MCSs in the subnetwork that knock out the function of the full network. Finally, like EFPs, MPs are defined for subnetworks as well as full networks, which allows one to adjust the subnetwork size for complete pathway enumeration in practice.

We argue that our iterative minimization approach based on LPs can replace the computationally hard direct minimization of a single MILP in the state of the art for pathway enumeration and sampling^18,22,25^. Running time as a function of subnetwork size, normalized by the size of the full network, was described well by a double exponential model for both approaches. However, our iterative approach had a constant that was an order of magnitude smaller than the direct approach, and it scaled with a much smaller base for exponentiation, allowing MPs to be found in networks of all sizes. More specifically, our graph-based approach completed more enumerations within one hour than other methods, and it yielded an order of magnitude more MPs in most incomplete enumerations. However, the number of MPs found decreased slightly with subnetwork size, likely due to the identification of maximal cliques, which is computationally hard and suffers from the increasing size of the graph with the number of MPs^56^. Here, we detected all maximal cliques each time the graph was updated (when the full subnetwork was used to find an MP), but future implementations could benefit from directly finding cliques with edges between reactions in the latest MP.

For randomized iterative minimization, absolute running time scaled with absolute subnetwork size, reflecting that it needs to solve a number of LP problems proportional to subnetwork size. This would also explain why direct minimization outperformed randomized iterative minimization in subnetworks large in absolute terms, but small relative to the full network. Note that no complete enumerations were registered by the randomized algorithm because we did not check for completeness; this would require integer variables and constraints at significant performance cost. We emphasize that the main bottleneck for enumeration is the BIP used to find cut sets, which slows down each iteration, especially as the number of known MPs and MCSs increases. Thus, a faster procedure for finding new cut sets would be a major contribution toward further scaling. As a general guideline for applying our methods to (sub)networks of interest, we therefore suggest that graph-based enumeration is used first to check if MPs and MCSs are enumerable in reasonable time. If not, the subnetwork size can be reduced, additional constraints can be applied, or MPs can be sampled randomly rather than enumerated. We also recommend network compression, as commonly applied before EFM computation^14^, to reduce the initial problem sizes as much as possible.

We demonstrated that random sampling and enumeration of MPs can yield biological insights through two example applications. First, random sampling of MPs in *E. coli* central carbon metabolism within a genome-scale network revealed the relative importance of reactions and enzyme-encoding genes in better agreement with experimental gene essentiality data than FBA. This approach is straight-forward and could be valuable for many applications, for example for the selection of knock-outs unlikely to disrupt growth or product formation in metabolic engineering strategies. Second, we enumerated all minimal sets of metabolite exchanges in a host-microbe model of a human cell interacting with six gut microbes. This showed that graph-based enumeration can be used to analyze some of the largest models considered to date, and our predictions included metabolite exchanges with experimental evidence and possible roles in gut homeostasis and human health. We note that future studies of gut microbiotas could use relaxed problem formulations to generate more MPs for potential metabolite exchanges—here, we aimed to identify minimal requirements for validation against the experimental literature. Overall, we argue that MPs open up new possibilities for detailed analysis of large-scale metabolic networks, and we expect them to produce new biological insights into uni- and multicellular metabolic systems.

## Methods

### Minimal pathways and cut sets

We start from a CBM with a flux vector **r** constrained by (1) and (2). We define the metabolic network 𝒩 = {1, 2, …, *n*} as the set of reaction indices in **r**, and we define a subnetwork 𝒮 and a pathway 𝒫 such that 𝒫 *⊆* 𝒮 *⊆* 𝒩. We further require 𝒫 to be a minimal pathway (MP), here defined as a minimal set of reactions in S that need to be active (have non-zero flux) in order to satisfy (1), (2), and any other linear constraints on 𝒩. In other words, 𝒫 is *support-minimal*: if the only active reactions in 𝒮 are 𝒫, constraints on 𝒩 will be violated if any reaction in 𝒫 is deactivated. We denote the set of known MPs as Π.

We define a cut set 𝒞 *⊆* ∪ Π as a set of reactions that intersects every 𝒫 ∈ Π. This means that deactivating all reactions in 𝒞 will deactivate all known MPs. We also require 𝒞 to be support-minimal such that removing any reaction from 𝒞 will leave an empty intersection with at least one 𝒮 ∈ Π. A minimal cut set (MCS) deactivates all MPs in 𝒫 and thereby disables flux in 𝒩 as a whole. We denote the set of known MCSs as Γ.

### Model preprocessing

We use flux variability analysis (FVA;^7^) to determine tight bounds 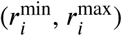 on feasible fluxes for all *i* ∈ 𝒩. Reactions in S are then made irreversible with positive upper bounds by replacing reversible reactions by two irreversible ones and reversing irreversible reactions with negative lower bounds. FVA also reveals blocked reactions,

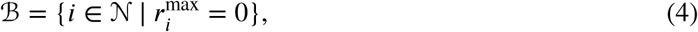

which always have zero flux and therefore are not part of any MP or MCS, and essential reactions,

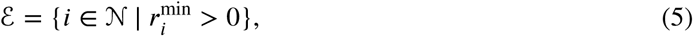

which cannot have zero flux and are MCSs that participate in all MPs. Hence, only the subnetwork 𝒮\(𝓑∪ 𝓔) can form MPs and MCSs along with 𝓔. For simplicity, we will keep denoting this subnetwork as 𝒮.

### Finding a new pathway with direct minimization

For each *i* ∈ 𝒩, we use a binary variable *b*_*i*_ ∈ {0, 1} to indicate if *r*_*i*_ is non-zero, as ensured by an inequality constraint:

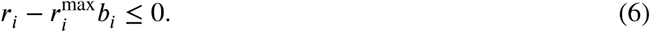

Another inequality constraint stops each 𝒫 ∈ Π from being found again:

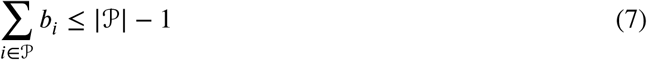

We can combine constraints (2), (6), and (7):

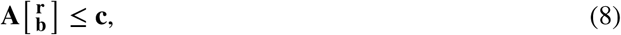

where the matrix **A** contains the coefficients, **b** is the vector of indicator variables, and **c** is the vector of constant terms. A new MP is then found by solving the MILP

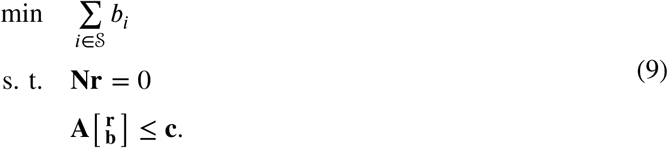

It minimizes the number of active reactions in S by minimizing the number of non-zero indicator variables. In an optimal solution, the active reactions in S define an MP that has the smallest possible number of active reactions among all MPs that have not yet been found.

### Finding a new cut set

We find a new cut set that deactivates all known MPs as described for EFMs by Song et al.^29^. Each reaction in at least one known MP, *i* ∈ ∪ Π, is represented by a binary variable *b*_*i*_ ∈ {0, 1}, and we use a constraint for each 𝒫 ∈ Π to ensure that it is covered by the cut set:

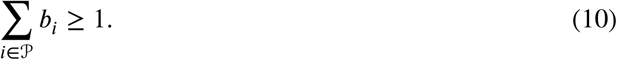

We also use a constraint for each 𝒞 ∈ Γ to exclude known MCSs:

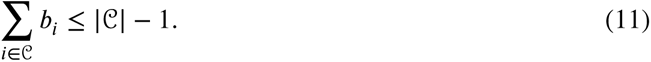

By reformulating constraints (10) and (11) as in (8), a new cut set results from solving the binary integer program (BIP)

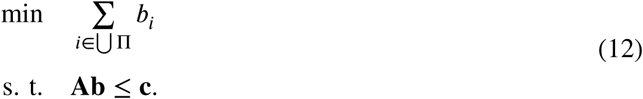

Specifically, a new cut set is given by the non-zero binary variables in the optimal solution; it may or may not be an MCS.

To find a new MP with iterative minimization, it is necessary to find a *viable* cut set that intersects all 𝒫 ∈ Π but does not intersect at least one unknown MP, i.e., a cut set that is not an MCS. To achieve this, **Algorithm S1** repeatedly solves and constrains the BIP and checks viability with the LP.

### Finding a new pathway with iterative minimization

Assuming that a cut set that disables known MPs has been applied, our iterative method (**Algorithm S2**) does not require integer variables or additional constraints to find a new MP. At each iteration, a reaction *i* ∈ 𝒮 is deactivated before minimizing the sum of fluxes in 𝒮 in the LP

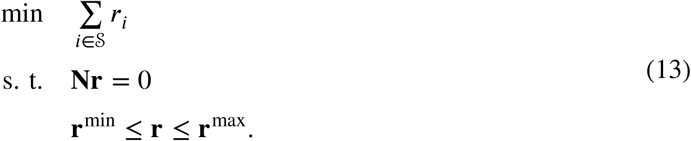

If an optimal solution is found, reactions in 𝒮 with zero flux in the solution can be deactivated as well because they are not needed in 𝒫. If no optimal solution is found, *i* must be part of 𝒫 and is reactivated. We repeat this until all *i* ∈ 𝒮 have been tested, and the remaining active reactions in 𝒮 constitute an MP that is support-minimal but does not necessarily have the smallest possible number of reactions.

We can also randomize the order of MPs by deactivating a random *i* ∈ 𝒮 at each iteration. In this case, we cannot deactivate all reactions in 𝒮 with zero flux in an optimal solution because randomness would not be preserved. However, random reactions can be drawn and deactivated until a reaction with non-zero flux in the optimal solution is drawn.

### Simple enumeration and sampling of pathways

Direct minimization can be used to enumerate MPs in order of increasing size by repeating the following steps until the MILP becomes infeasible:

1. Find a new MP by solving the MILP.
2. Add constraint (7) to the MILP to avoid finding the MP again.

Iterative minimization can be combined with the BIP for finding new cut sets to enumerate MPs based on the procedure by Song et al.^29^. The following steps are repeated until the BIP becomes infeasible:

1. Find and apply a new viable cut set using **Algorithm S1**.
2. Find a new MP using **Algorithm S2**.
3. Add constraint (10) to the BIP to avoid finding the cut set again.

This produces all MPs (not in order of increasing size) as well as all MCSs.

If too many MPs exist for complete enumeration to be feasible, MPs can be sampled randomly by repeatedly finding random MPs without finding and applying cut sets until the number of MPs equals the desired sample size. This avoids the significant overhead of solving BIPs but risks finding the same MP multiple times if the number of MPs is not sufficiently large.

### Graph-based enumeration of pathways

An undirected graph can be used to predict new MPs from known MPs. We define this graph as the union of reaction pairs from all 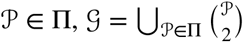, where reactions are nodes and reaction pairs are edges connecting two nodes. Reactions are connected if they occur together in at least one known MP, meaning that MPs are *cliques* (complete subgraphs) in G. We exploit that edges defined by known MPs can combine with each other to form additional cliques that may contain previously unknown MPs.

Each time we find a new MP during enumeration, we update the graph by adding any missing edges between reactions in the new MP. We use a modified Bron-Kerbosch algorithm^56^ to find *maximal cliques*, i.e., fully connected subgraphs where no other nodes in the graph are connected to all nodes in the subgraph. The MPs in each new maximal clique are enumerated separately using iterative minimization with the clique as subnetwork in place of 𝒮. This reduces the sizes of the optimization problems and thereby accelerates enumeration.

Graph-based prediction of new MPs can be combined with direct or iterative minimization (or any other procedure that returns new MPs); here, we implemented iterative minimization as described in **Algorithm S3**. However, graph-based prediction is biased by the known MPs and therefore incompatible with random sampling of MPs.

### Benchmarking

We obtained six CBMs (Table S1) from the BiGG database^2^, comprising models of *Escherichia coli*^30,57^, *Helicobacter pylori*^58^, *Saccharomyces cerevisiae*^59^, *Cricetulus griseus*^60^, and *Homo sapiens*^61^. We removed any non-growth-associated maintenance reactions and all biomass reactions except the one with the largest number of metabolites, set the default bounds of fluxes to ± 1,000 mmol gDW^−1^ h^−1^, set a minimal growth rate of 0.1 h^−1^, and allowed uptake and secretion of all environmental metabolites to simulate a complex medium. We preprocessed each model as described above.

For each model, we sampled random subnetworks ranging from ten reactions to the full network. We chose logarithmically spaced subnetwork sizes with four sizes sampled for each order of magnitude between 10 and 10^4^. We sampled 100 random subnetworks of each size and enumerated MPs using direct and iterative minimization (with and without graph or randomization). Incomplete enumerations were stopped after one hour.

We used bootstrapping to compute 95% confidence intervals for estimated means. The data were resampled 1,000 times with replacement, the mean was reestimated for each sample, and a 95% confidence interval was defined from the 2.5th and 97.5th percentiles of these estimates.

### Application to central metabolism

We obtained a pathway map of *E. coli* central carbon metabolism from Escher^62^ and used the reactions in this map as a subnetwork within the genome-scale model iJO1366^36^. Using randomized iterative minimization, we sampled 100,000 MPs in the subnetwork with a growth requirement of 0.1 h^−1^, the default minimal glucose medium, and aerobic conditions. We defined reaction (gene) importance as the fraction of sampled MPs requiring each reaction (enzyme-encoding gene). To map samples from reactions to genes, we computed the contribution of each gene to each reaction by dividing the number of enzymes requiring the gene by the total number of enzymes catalyzing the reaction. To predict gene importance with FBA, we deleted each gene from the model, maximized growth rate, and computed the fractional deviation from the maximal growth rate of the wild type. We obtained growth and gene essentiality data from the Keio collection^32^. Bootstrapping was performed as for benchmarking.

### Application to host-microbe exchanges

We built a host-microbe model in which genome-scale CBMs of a human cell and of six commensal gut microbes (*Bacteroides thetaiotaomicron*, *E. coli*, *Faecalibacterium prausnitzii*, *Lactobacillus plantarum*, *Lactococcus lactis*, and *Streptococcus thermophilus*; Table S2) can exchange metabolites through the gut lumen. We based the model on previous work^63^ but used a more recent human model^61^ and collection of gut microbe models^43^, all obtained from Virtual Metabolic Human^64^. We set a non-growth-associated ATP requirement of 0.1 mmol gDW^*1^ h^*1^ for the human cell and a growth requirement of 0.1 h^*1^ for the microbes. To simulate the anaerobic and potentially complex metabolic environment of the gut, we allowed uptake and secretion of all environmental metabolites for all cells, turned off oxygen exchange for the microbes, and added reactions representing all possible metabolite exchanges between microbes and from microbes to the human cell (excluding ATP, sugars, and inorganic ions and compounds). Building on our recent approach^42^, we also applied two FBA-steps, minimizing (1) microbial intracellular flux^53^ and (2) uptake flux not explained by metabolite exchange across all cells:

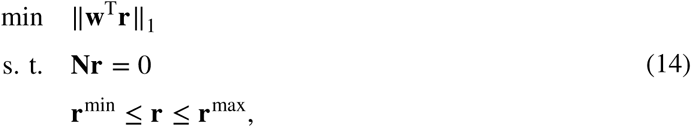

where ||**w**^T^**r**||_1_ is the weighted sum of absolute fluxes to be minimized. After the first step, we constrained each intracellular microbial flux to its value in the minimum obtained. After the second step, we constrained the absolute sum of exchange fluxes not explained by metabolite exchange to the minimum obtained, 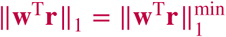. This approach aimed to explain as much of the functional requirements as possible in terms of metabolite cross-feeding with biologically reasonable microbial flux distributions. We enumerated minimal sets of metabolite exchanges using iterative minimization with graph.

### Software

We implemented the algorithms in Python using COBRApy^65^ and the Gurobi Optimizer (Gurobi Optimization, LLC, Beaverton, OR, USA).

## Data and code availability

The data and code needed to reproduce our results are available at https://gitlab.com/csb.ethz/mptool.

## Acknowledgements

This work was supported by the SystemsX.ch project GutX, evaluated by the Swiss National Science Foundation, and the Swiss National Science Foundation Sinergia project #177164. We thank Mattia Gollub, Axel Theorell, and the DigiSal team at the Norwegian University of Life Sciences for helpful comments on the manuscript.

## Author contributions

J.S. and O.Ø. designed the study and wrote the manuscript. O.Ø. developed the algorithms and performed the analyses.

## Supplementary information

**Table S1.** Models used for benchmarking with number of metabolites and reactions before (and after) preprocessing.

| Organism | Model | Reactions | Metabolites |
| --- | --- | --- | --- |
| *E. coli* | e_coli_core | 95 (137) | 72 (72) |
| *H. pylori* | iIT341 | 554 (600) | 485 (422) |
| *S. cerevisiae* | iMM904 | 1,577 (1,273) | 1,226 (722) |
| *E. coli* | iML1515 | 2,712 (2,868) | 1,877 (1,633) |
| *C. griseus* | iCHOv1 | 6,663 (6,805) | 4,456 (2,749) |
| *H. sapiens* | Recon3D | 10,600 (14,119) | 5,835 (5,835) |

**Table S2.**
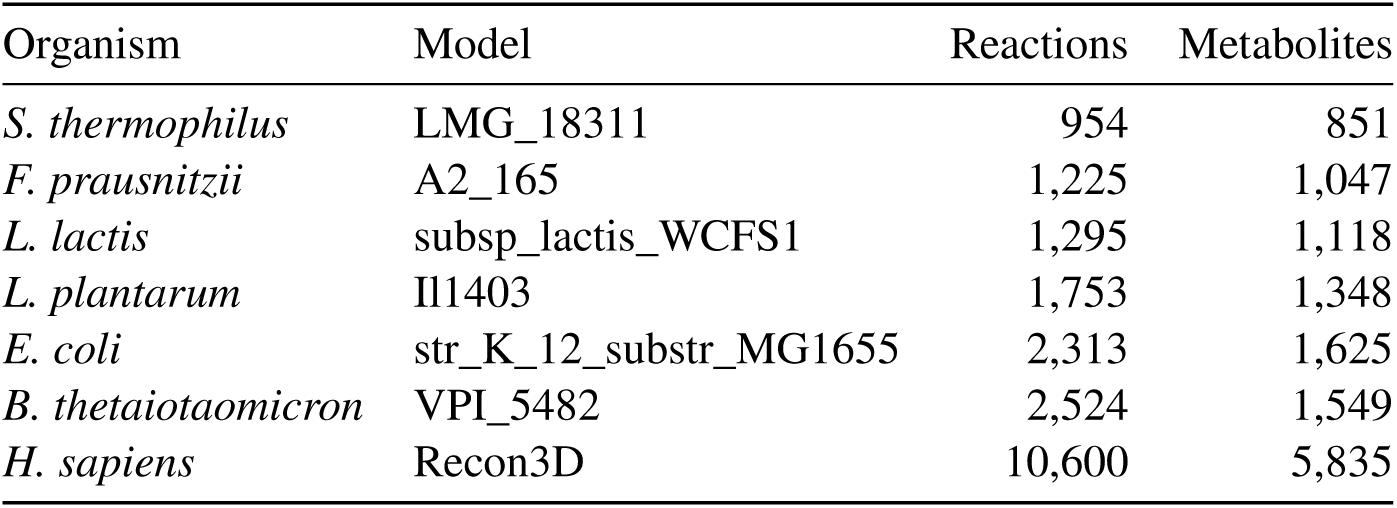
Models used in host-microbe model with number of metabolites and reactions. Identifiers of microbial models are from Magnúsdóttir et al. ^43^.

### Algorithm S1: Finding a new viable cut set

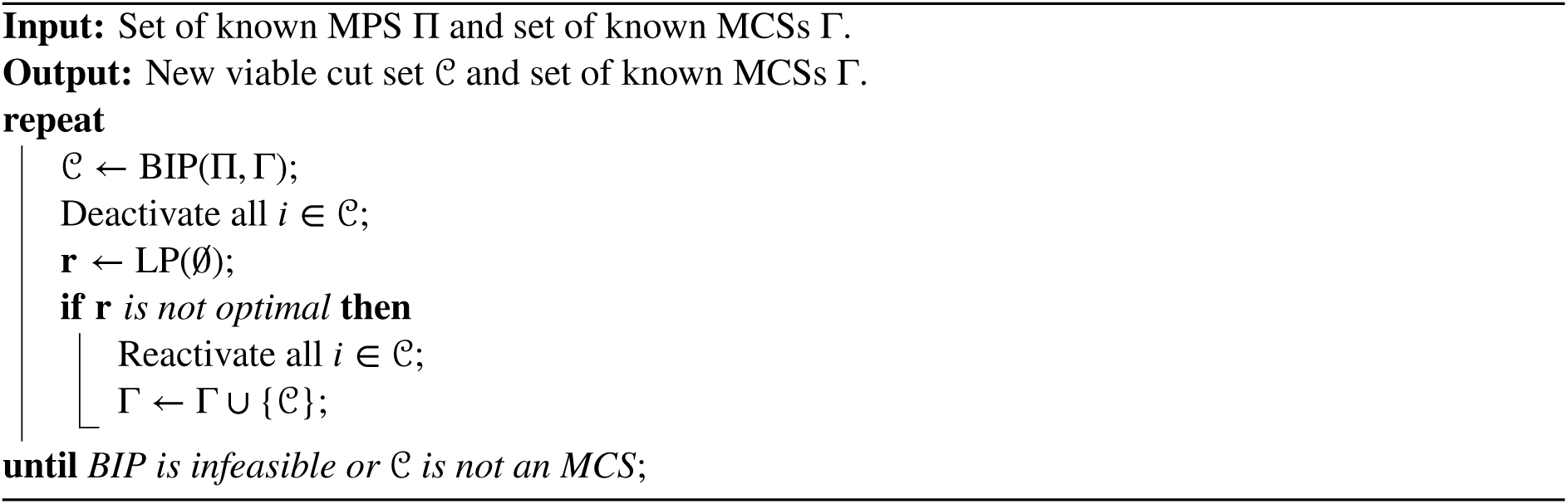

### Algorithm S2: Finding a new MP with iterative minimization

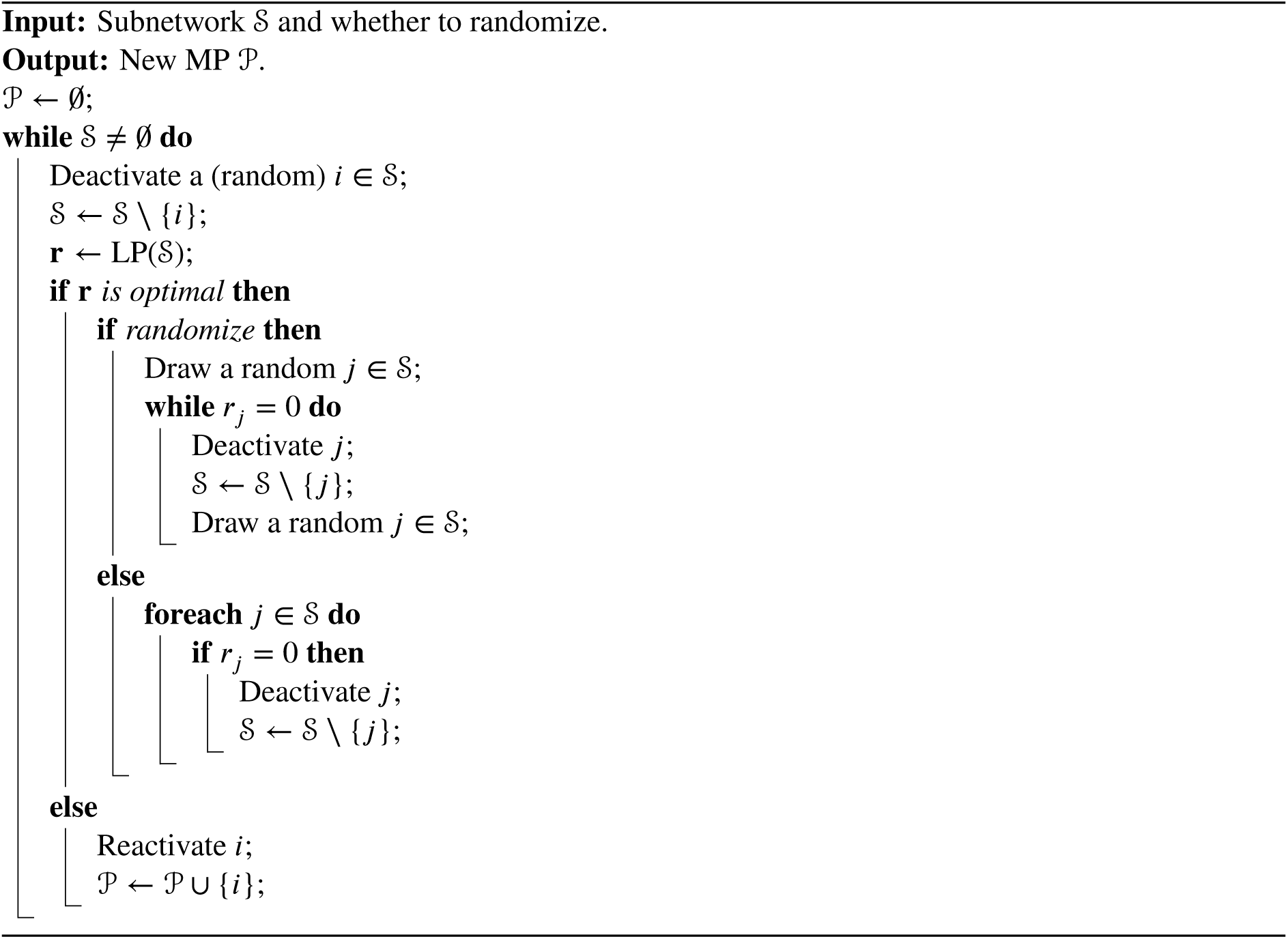

### Algorithm S3: Enumeration with iterative minimization and graph

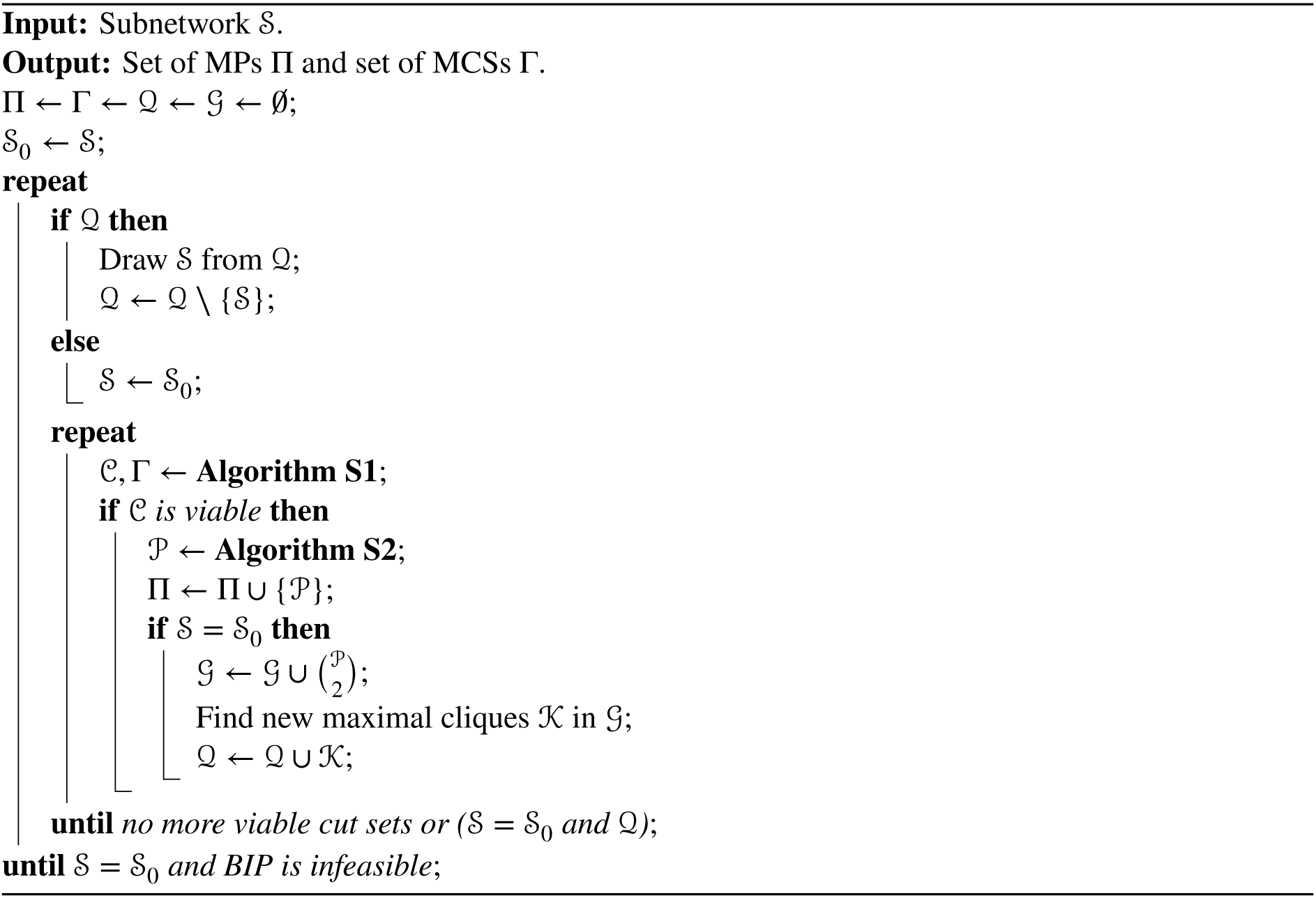

**Figure S1.**
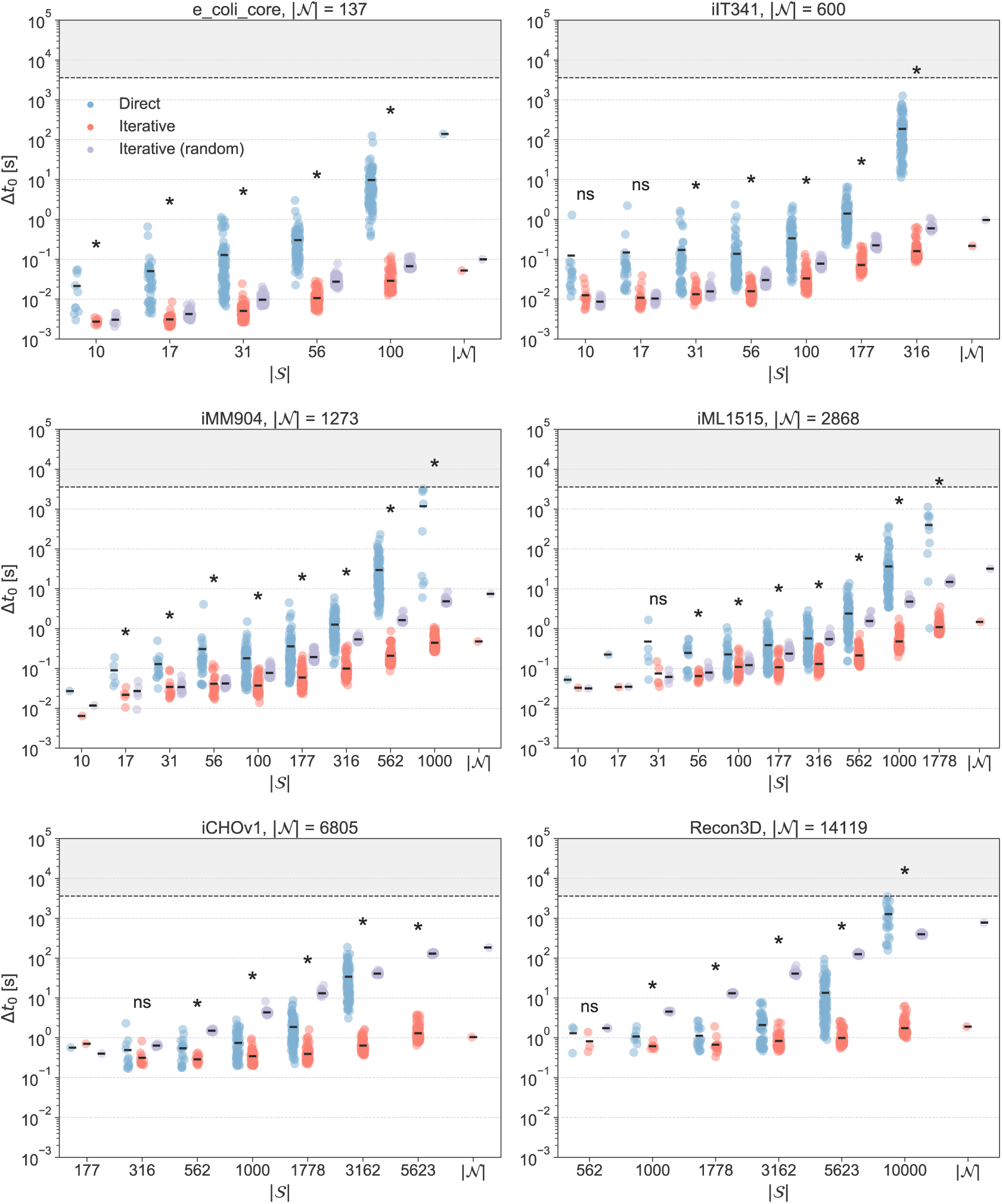
Running time *t*_0_ for finding the first new MP with direct (blue) and iterative minimization with (purple) and without (red) randomization for different subnetwork sizes |𝒮| in six different models with network size |𝒩|. Stars indicate significant difference between means from one-way ANOVA where data was available for all three methods or *t*-test where data was available for two methods (significance level 0.05). Optimization was stopped after one hour (indicated by dashed line).

**Figure S2.**
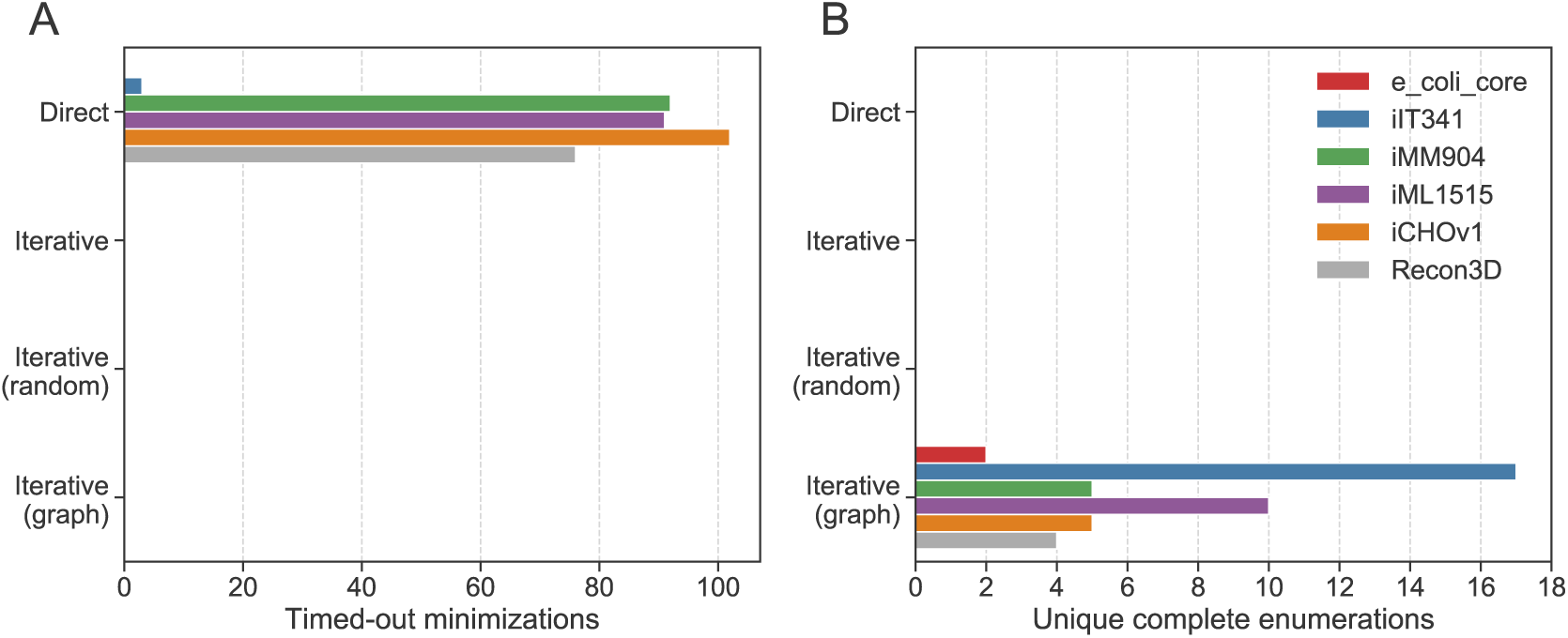
(**A**) Number of minimizations that timed out before the first MP was found and (**B**) number of enumerations completed by one method but not by any of the others by method and model.

**Figure S3.**
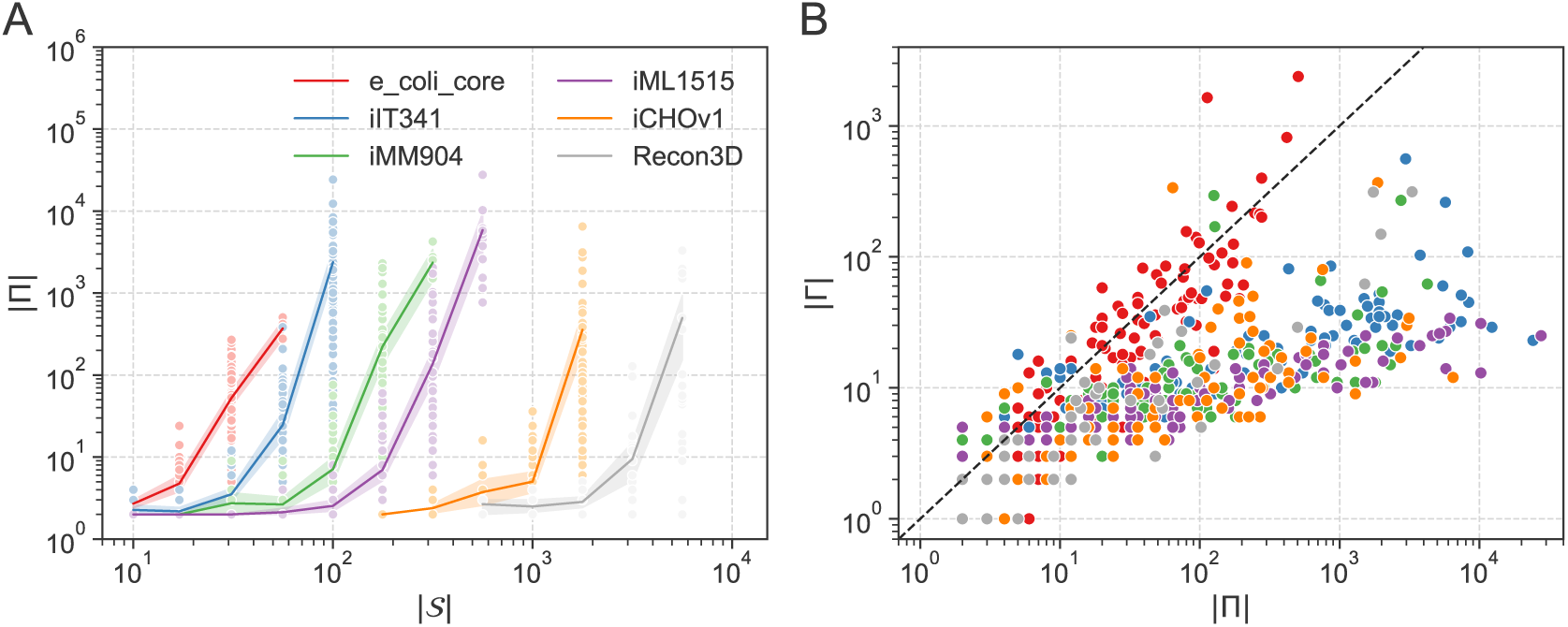
MPs found in complete enumerations. (**A**) Number of MPs |Π| as a function of subnetwork size |𝒮| for enumerations completed within one hour by at least one method. Lines indicate means with 95% confidence intervals from bootstrapping and colors indicate models. (**B**) Number of MCSs |Γ| as a function of |Π| with colors indicating models and a dashed line indicating |Γ| = |Π|.

**Figure S4.**
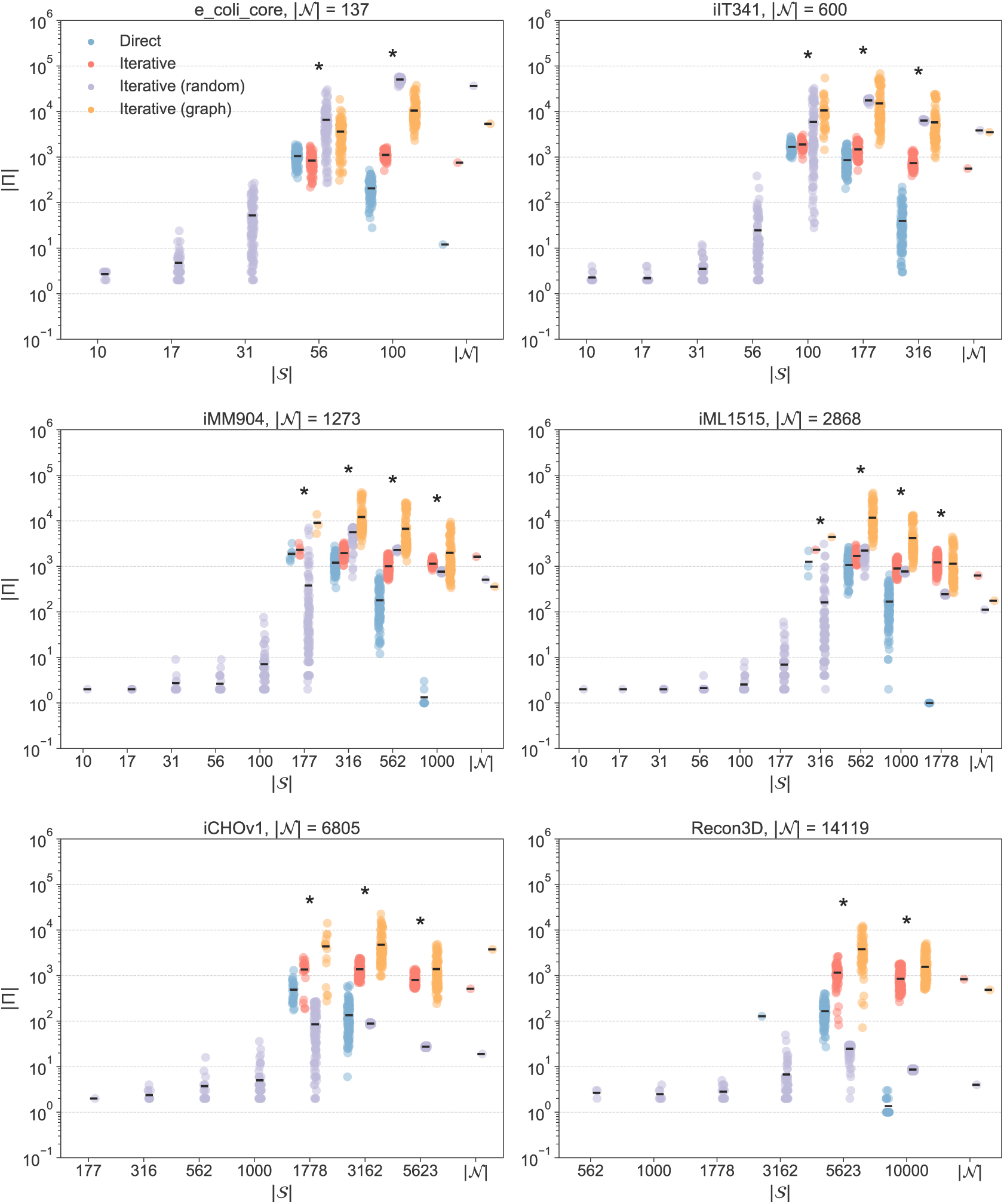
Number of MPs |Π| found in one hour with direct minimization (red), iterative minimization (blue), iterative minimization with randomization (purple), and iterative minimization with graph (orange) for different subnetwork sizes |𝒮| in six different models with network size |𝒩|. Stars indicate significant difference between means from one-way ANOVA where data was available for all three methods or *t*-test where data was available for two methods (significance level 0.05). Enumerations that were completed within one hour are not included.

**Figure S5.**
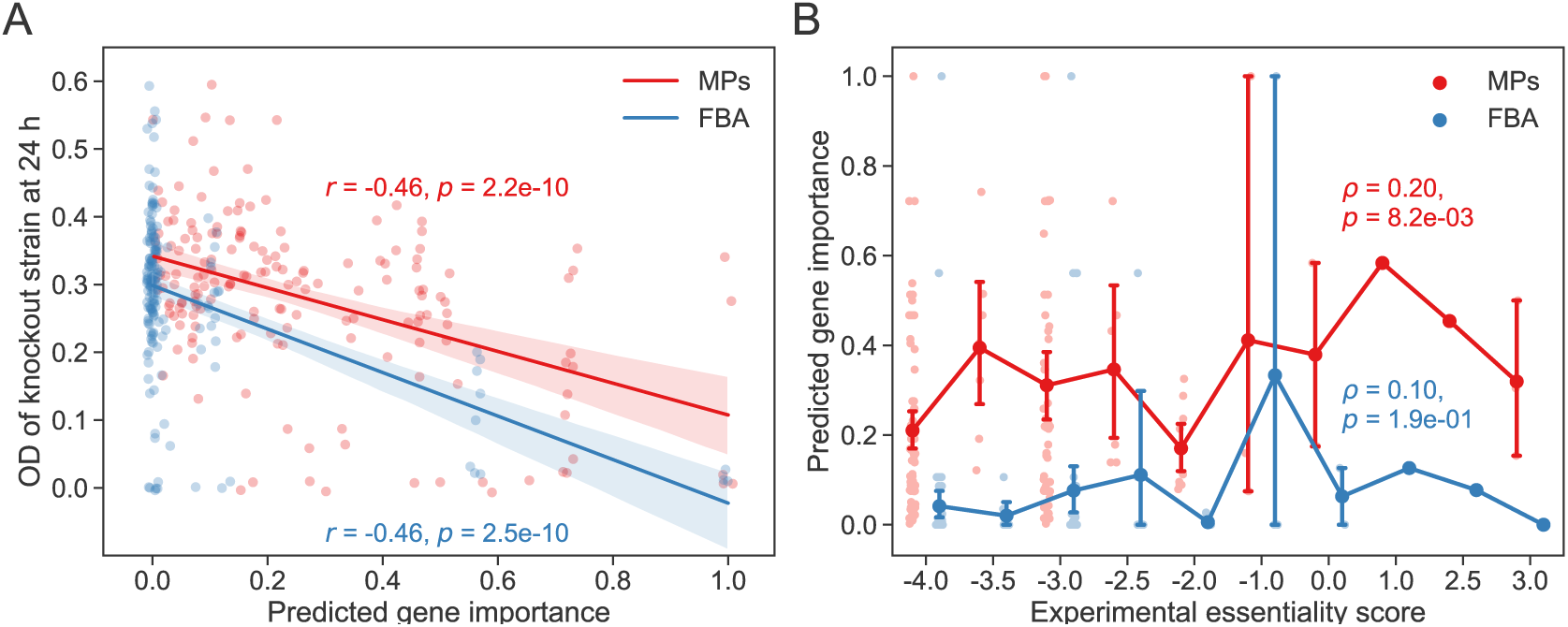
Correlation of gene importance in *E. coli* central carbon metabolism predicted by MP sampling (red) and FBA (blue) with experimental growth and gene essentiality data from the Keio collection ^32^. (**A**) Pearson correlation (*r*) of predicted gene importance with experimental growth (OD of knockout strain at 24 h). Fitted lines are shown with 95% confidence intervals from bootstrapping. (**B**) Spearman correlation (*ρ*) of predicted gene importance with experimental gene essentiality score (higher score means more evidence for essentiality and vice versa). Means are indicated with 95% confidence intervals from bootstrapping.

